# Isolation of bacteria that catabolize abiotically synthesized tetroses via the formose reaction

**DOI:** 10.1101/2024.11.09.622817

**Authors:** Kosei Kawasaki, Kensuke Igarashi, Hiro Tabata, Hiroaki Nishijima, Shuji Nakanishi, Souichiro Kato

## Abstract

Sugars synthesized through abiotic processes, specifically the formose reaction, are promising candidates for next-generation feedstocks for biomanufacturing due to their significantly higher productivity compared to cultivated crops. One of the significant challenges associated with the utilization of abiotically synthesized sugars via the formose reaction is the presence of unusual sugars, which are not metabolized by typical microorganisms. This study aimed to identify microorganisms capable of catabolizing tetroses, which are among the unusual sugars present in the abiotically synthesized sugars. Four model bacteria commonly used in biomanufacturing, including *Escherichia coli*, were unable to catabolize tetroses and their growth was completely inhibited by 2 to 4 g/L of D-erythrose or 0.5 to 1 g/L of L-erythrulose. We successfully isolated eight phylogenetically diverse bacterial strains from river sediments, capable of utilizing D-erythrose and L-erythrulose as the sole carbon sources. These isolates exhibited higher tolerances to tetroses compared to the model bacteria. The isolates exhibited the ability to grow in high concentrations of the abiotically synthesized sugars, which inhibit the growth of typical microorganisms, and to consume tetroses contained therein. Further investigation into the molecular mechanisms of tetrose metabolisms in the isolates will facilitate the development of biomanufacturing processes utilizing the abiotically synthesized sugars.

## INTRODUCTION

The technology for synthesizing valuable compounds using biological systems, known as biomanufacturing, has garnered significant attention due to its energy efficiency, reduced dependence on fossil resources, and its pivotal role in fostering a sustainable economy^1,2^. Conventional biomanufacturing predominantly utilizes sugars derived from cultivated crops as feedstocks. However, concerns regarding competition with food, land use issues, and the depletion of water resources have necessitated the development of more sustainable feedstocks^3,4^. In such contexts, research has focused on the use of inedible biomass^5^ and algal biomass^6^ as the alternative feedstocks, as well as on the direct production of organic compounds from CO_2_ using non-photosynthetic microorganisms such as hydrogen-oxidizing bacteria, acetogenic bacteria, and electrosynthetic bacteria^7-14^. The utilization of organic compounds such as acetic acid, which can be rapidly synthesized through electrocatalytic CO_2_ reduction, as feedstocks for biomanufacturing is also considered a promising technology^15,16^.

In recent years, sugars synthesized through abiotic processes, specifically the formose reaction, have attracted interest as promising alternative feedstocks for biomanufacturing^17-19^. The formose reaction is known as a sugar formation reaction, primarily involves heating formaldehyde in an alkaline solution in the presence of a metal catalyst such as calcium hydroxide. Since its discovery in 1861, the formose reaction has been widely studied as a source of sugars in the origin of life^20-22^. Given that the rate of sugar synthesis via the formose reaction significantly surpasses that of cultivated crops, several research groups explored the application of the abiotically synthesized sugars in biomanufacturing during the 1960s and 1970s^23,24^. However, the formose reaction is a highly complex process with numerous undesired reactions, leading to the formation of an extremely diverse sugars and their derivatives. Due to this property, the bioavailability of the abiotically synthesized sugars is extremely low, which has hindered their practical application.

Our research group has discovered that metal oxoacids, such as sodium tungstate and sodium molybdate, function as catalysts for the formose reaction under neutral conditions^25^. These conditions minimize undesired reactions, thereby facilitating efficient sugar synthesis with relatively high selectivity. In addition, we demonstrated that the abiotically synthesized sugars via the formose reaction functions as growth substrates for model biomanufacturing bacteria such as *Escherichia coli* and *Corynebacterium glutamicum*^19^. These properties suggest that the abiotically synthesized sugars could serve as alternative feedstocks for biomanufacturing through further improvements in the formose reaction. One of the significant challenges associated with the abiotically synthesized sugars is the presence of unusual sugars, such as branched sugars and L-form sugars, which are not metabolized by typical microorganisms. Nevertheless, we demonstrated that a significant portion of the abiotically synthesized sugars is consumed by soil microbial communities^25^. This finding suggests that microorganisms capable of catabolizing unusual sugars are present in natural environments and that the bioavailability of the abiotically synthesized sugars can be enhanced by leveraging their metabolic abilities.

Tetroses are among the unusual sugars present in the abiotically synthesized sugars. Tetroses are four-carbon sugars with six isomers, comprising two aldoses (erythrose [ery] and threose) and one ketose (erythrulose [eru]), each of which has both D and L configurations. In the previous study, we showed that tetroses contained in the abiotically synthesized sugars inhibit the growth of *C. glutamicum*^19^. However, the understanding of microbial metabolism and the biotoxicity of tetroses remains extremely limited, likely due to the rarity of tetroses in natural environments. In order to utilize the abiotically synthesized sugars for biomanufacturing, it is essential to investigate hitherto unknown microorganisms capable of metabolizing and tolerating tetroses.

In this study, we first investigated the tetroses availability and sensitivity of model bacteria commonly used for biomanufacturing. We then attempted to isolate microorganisms capable of growing under conditions where tetroses serve as the sole carbon and energy source. The obtained isolates were examined for their tetrose catabolizing abilities and tolerance properties, as well as their capacity to grow on the abiotically synthesized sugars and consume the tetroses contained therein.

## MATERIALS AND METHODS

### Bacterial Strains and Culture Conditions

*E. coli* strain K12 (ATCC 12435) was routinely cultured in the M9 minimal medium^26^ supplemented with 1 g/L D-glucose (D-glc). *Bacillus subtilis* strain 168 (JCM 10629) and *C. glutamicum* strain 534 (ATCC 13032) were regularly cultured in the IET minimal medium^27^ supplemented with 1 g/L of D-glc. *Cupriavidus necator* (DSM 428, formerly known as *Ralstonia eutropha*) was routinely cultured in the IET minimal medium supplemented with 1 g/L of D-fructose (D-fru). All model strains were cultured at 30°C with agitation (180 rpm). D-ery and L-eru were obtained from Sigma-Aldrich Japan (Tokyo, Japan). Stock solutions of each sugar were prepared by filter sterilization using 0.22-μm pore filters (MilliporeSigma, Burlington, MA, US) and subsequently added to the autoclaved minimal media. Bacterial growth was assessed by measuring the optical density at 600 nm (OD_600_) of the culture solution with a 96-well plate reader (BioTek LogPhase 600, Agilent Technologies, Santa Clara, USA) using the uninoculated medium as the control.

### Isolation of Tetrose-Utilizing Microorganisms

Tetrose-utilizing microorganisms were enriched in test tubes filled with 1 ml of the IET minimal medium supplemented with 2 g/L of D-ery or L-eru and 5 μg/mL of the fungal inhibitor amphotericin B. Approximately 50 μL of river sediment collected from Sakuskotoni and Atsubetsu river (Sapporo, Japan) was inoculated as a source of microorganisms. Enrichment cultures were incubated at 30°C with agitation (180 rpm) until visible growth was confirmed (2 to 4 days). After at least three successive enrichments, the serially diluted culture solution was inoculated onto the agar-solidified IET minimal medium supplemented with 10 g/L D-glc to obtain isolates by single colony isolation. The isolates were further purified by single colony isolation on the same medium at least twice. Subsequently, the ability of the isolates to utilize D-ery or L-eru was assessed using the IET liquid medium to identify tetrose-utilizing strains. All tetrose-utilizing isolates were routinely cultured in the IET minimal media supplemented with 1 g/L D-glc at 30°C with agitation (180 rpm). Almost the full length of the 16S rRNA gene sequences of the isolates were determined by the direct sequencing of PCR products with the primer pair 27F/1492R as described previously^28^. Isolates with >98% 16S rRNA gene sequence identities were designated as same phylotypes, and one representative strain from each phylotype was subjected for further analysis. The closest relatives of the isolates were inferred using the BLAST program^29^.

### Evaluation of Utilization of the Abiotically Synthesized Sugars by Isolates

The tetrose-utilizing isolates were cultured in the IET minimal medium supplemented with 5% (w/v) of the abiotically synthesized sugars prepared as described previously^19^. Formaldehyde (C1), glycolaldehyde (C2), glyceraldehyde (C3a), dihydroxyacetone (C3k) and tetroses (C4a and C4k) (in this study, substances with n carbons are denoted as Cn, with aldoses and ketoses denoted as Cna and Cnk, respectively) in the supernatant of culture solution were quantified using a high-performance liquid chromatography (Chromaster system, Hitachi High-Tech, Tokyo, Japan) as described elsewhere^19^.

## RESULTS

### Inhibitory Effects of Tetroses on Model Bacteria

The growth capabilities and resistance properties of model bacteria to tetroses were investigated. The wild types of *E. coli, C. necator, B. subtilis*, and *C. glutamicum* were used as the model bacterial strains commonly used in biomanufacturing. D-ery and L-eru were used as the representatives for C4a and C4k, respectively.

Each model strain was cultured in the minimal media supplemented with different concentrations (1, 2, or 4 g/L, corresponding to 8.3, 16.7 or 33.3 mM) of D-ery and L-eru. No significant growth was observed in any of the cultures (data not shown), indicating that these bacteria are incapable of utilizing tetroses as the sole carbon and energy source. In order to evaluate the growth-inhibitory effects of tetroses, the model strains were cultured in the minimal media supplemented with 1 g/L D-glc (or D-fru for *C. necator*) and varying concentrations (0.5, 1, 2, or 4 g/L) of D-ery or L-eru. Despite some differences between strains, the four model strains showed similar trends, *i*.*e*., their growth was completely inhibited by 2 to 4 g/l of D-ery or 0.5 to 1 g/L of L-eru (Figure 1). These results indicate that tetroses, especially C4k, have significant inhibitory effects on bacterial growth. It should be noted that lower concentrations of tetroses promoted the growth of *C. glutamicum* (Figure 1D). This suggests that *C. glutamicum* may possess the ability to metabolize tetroses as a supplementary carbon source, although tetroses cannot serve as the sole carbon or energy source.

**Figure 1.**
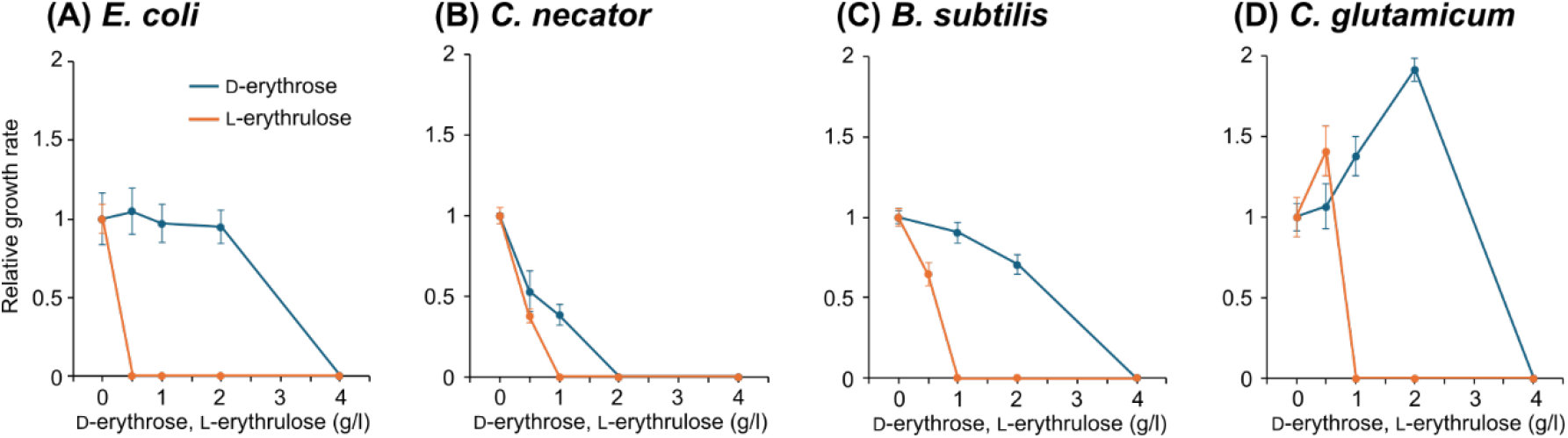
The inhibitory effects of D-erythrose and L-erythrulose on the growth of model bacterial species. The specific growth rates (*μ*, hr^-1^) were determined using the minimal media containing 1 g/L of D-glucose (or D-fructose for *C. necator*) and varying concentrations (0.5, 1, 2, or 4 g/L) of D-erythrose or L-erythrulose. The relative growth rates were determined by comparing the specific growth rates in these media to that in the glucose-only medium. Data are presented as the means of five independent cultures. Error bars represent standard deviations.

### Isolation of Tetrose-Utilizing Microorganisms

We aimed to isolate microorganisms capable of catabolizing tetroses, which are essential for employing the abiotically synthesized sugars as biomanufacturing feedstocks. Isolation of microorganisms was performed using the IET minimal medium with 2 g/L of D-ery or L-eru as the sole carbon and energy source and river sediments as the microbial sources. We successfully obtained five and three phylogenetically unique isolates capable of growing on D-ery and L-eru, respectively (Table 1). Phylogenetic analysis based on 16S rRNA gene sequences showed that the tetrose-utilizing isolates belong to a diverse array of lineages, *i*.*e*., Actinomycetota and α-, β-, and γ-proteobacteria, indicating that the ability to catabolize tetroses is not confined to a specific bacterial lineage. Of the eight isolates, four strains that grew well on D-ery or L-eru (strains Ery-7, Ery-28b, Eru-24, and Eru-33a) were subjected for further analysis.

**Table 1.** Phylogenetic information of the tetrose-utilizing isolates obtained in this study.

| Strain | Phylum/Class | Closest relatives (accession numbers) [identity, %] |
| --- | --- | --- |
| Isolates from D-erythrose enrichments |  |  |
| Ery-7 | γ-proteobacteria | *Serratia marcescens* AFS044658 (OP986802) [99.8] |
| Ery-15 | α-proteobacteria | *Brucella pseudogrignonensis* ESL2 (CP091784) [99.9] |
| Ery-23 | γ-proteobacteria | *Pseudomonas protegens* E2HL9 (MK855122) [99.9] |
| Ery-27 | β-proteobacteria | *Delftia acidovorans* FDAARGOS_939 (CP065627) [100] |
| Ery-28b | Actinomycetota | *Curtobacterium citreum* AFS070301 (OP986288) [99.9] |
| Isolates from L-erythrulose enrichments |  |  |
| Eru-6 | Actinomycetota | *Curtobacterium citreum* PgBE223 (MH144294) [99.8] |
| Eru-24 | β-proteobacteria | *Paraburkholderia humisilvae* MAW7 (OR759236) [98.6] |
| Eru-33a | β-proteobacteria | *Paraburkholderia aromaticivorans* BN5 (CP022989) [99.8] |

### Growth Capabilities and Resistance Properties of the Isolates to Tetroses

Four isolated strains were cultured in the minimal media supplemented with different concentrations (1, 2, or 4 g/L) of D-ery or L-eru to evaluate their tetrose-catabolizing capabilities. Strains Ery-7 and Ery-28b, isolated as D-ery-utilizing bacteria, exhibited growth at concentrations of 1 and 2 g/L of D-ery, but not at 4 g/L (Figure 2A and B). These strains did not exhibit significant growth on L-eru, while slight OD increases were observed in the media supplemented with 1 or 2 g/L of L-eru (Figure 2E and F). Strains Eru-24 and Eru-33a, isolated as L-eru-utilizing bacteria, were able to grow on both D-ery and L-eru. Strain Eru-24 exhibited growth at concentrations of 1 and 2 g/L of D-ery and L-eru, but not at 4 g/L (Figure 2C and G). In contrast, strain Eru-33a was able to grow on 4 g/L of D-ery and L-eru, although some growth retardation was observed (Figure D and H). These results unequivocally demonstrate that these isolates can utilize tetroses as their sole carbon and energy source. These results also indicate that, although bacteria possess the ability to catabolize tetroses, their growth is inhibited by high concentrations of these sugars.

**Figure 2.**
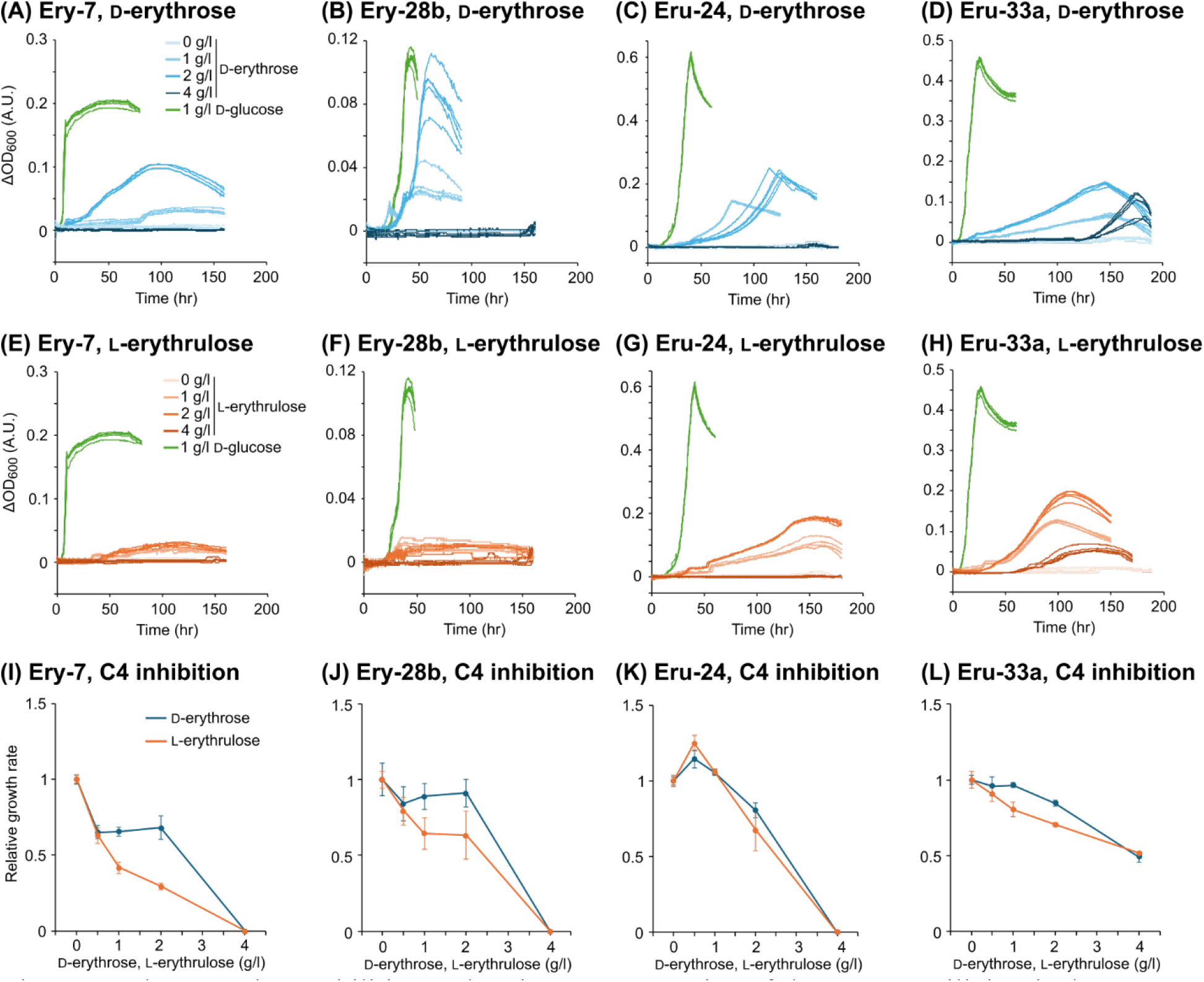
The growth capabilities and resistance properties of the tetrose-utilizing isolates on D-erythrose and L-erythrulose. (A-D and E-H) The growth curves of each isolate on different concentrations of D-erythrose and L-erythrulose, respectively. All data from five independent cultures are shown. As a comparison, growth curves on the media supplemented with 1 g/L D-glucose are shown (green lines). (I-L) The inhibitory effects of D-erythrose and L-erythrulose on the growth of the tetrose-utilizing isolates. The relative growth rates were determined as noted in Figure 1. Data are presented as the means of five independent cultures. Error bars represent standard deviations.

The isolates were cultured in the minimal media supplemented with 1 g/L of D-glc and varying concentrations (0.5, 1, 2, or 4 g/L) of D-ery or L-eru to evaluate their tolerance to tetroses. Despite some differences among the strains, strains Ery-7, Ery-28b and Eru-24 exhibited similar trends, *i*.*e*., their growth was completely inhibited by 4 g/L of D-ery or L-eru (Figure 2I-K). On the other hand, strain Eru-33a was able to grow in the media supplemented with 4 g/L of D-ery or L-eru, albeit with some growth suppression (Figure 2L). All of the tetrose-utilizing isolates exhibited higher resistance, particularly to L-eru, compared to the model bacteria.

### Utilization of the Abiotically Synthesized Sugars by Isolates

Four isolates were cultured in the minimal medium containing the abiotically synthesized sugars at concentrations that significantly inhibited the growth of the model strains (5% w/v) ^19^. Since the growth of all isolates was observed (data not shown), the C4a and C4k in the culture media were quantified over time (Figure 3). The control cultures without bacteria exhibited no decrease in tetroses, while a slight increasing trend was observed, likely due to the evaporation of the medium (Figure S1A). Consumption of C4a was observed in the cultures of all isolates, with concentrations decreasing from 27.8 mM to 12.6 to 18.8 mM. This result is reasonable since all isolates have the ability to catabolize D-ery, one of the isomers of C4a. About half of C4a remained unconsumed, presumably because isomers other than D-ery were not utilized. As for C4k, significant consumption was observed exclusively in the strain Eru-33a culture (Figure 3D), which is reasonable given that strain Eru-33a exhibited better growth on L-eru compared to other isolates (Figure 2H). Conversely, it is intriguing that C4k was not consumed in the cultures of another L-eru-utilizing isolate, strain Eru-24 (Figure 3C). Strain Eru-24 might grow by consuming the readily available sugars other than L-eru contained in the abiotically synthesized sugars. The concentrations of C1 (formaldehyde, the raw material of the formose reaction) and the intermediates C2, C3a, and C3k in the culture solution were also quantified (Figure S1B and S2). Among the isolates, only strain Eru-33a almost completely consumed the C1-3 compounds (Figure S2D), indicating that this strain is highly efficient in utilizing not only tetroses but also shorter chain sugars.

**Figure 3.**
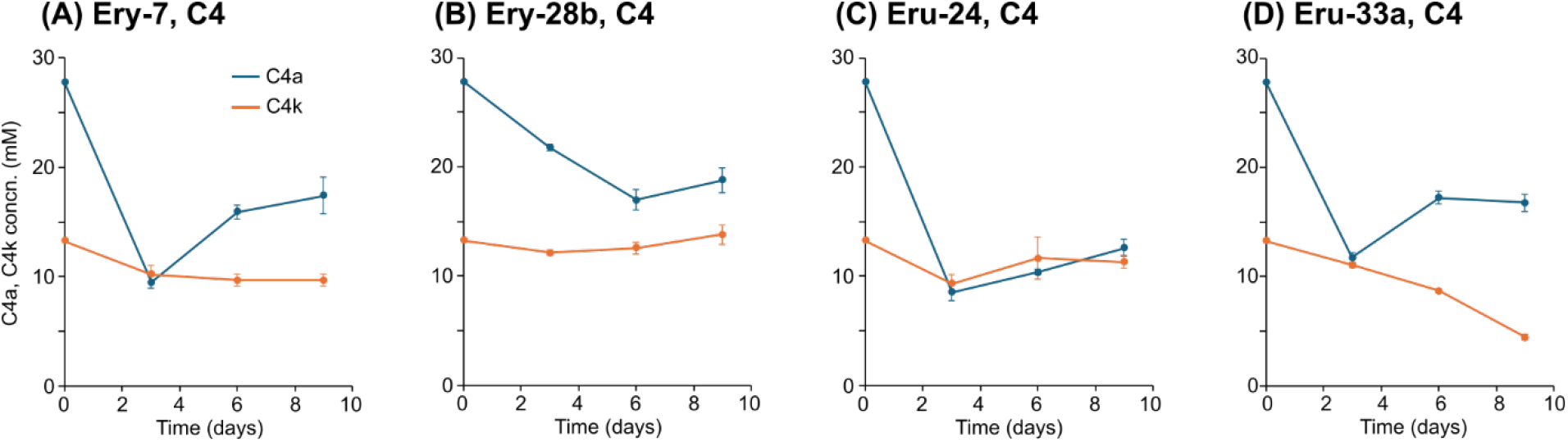
Consumption of C4 sugars in the abiotically synthesized sugars by the tetrose-utilizing isolates. Each isolate was cultured in the minimal medium supplemented with 5% of the abiotically synthesized sugars and the changes in the concentrations of each sugar species were determined. Data are presented as the means of three independent cultures. Error bars represent standard deviations.

## DISCUSSION

In this study, we clearly demonstrated that tetroses exhibit biotoxicity against phylogenetically diverse bacteria (Figure 1). Aldose and ketose sugars have been reported to exhibit biotoxicity due to the reactivity of their aldehyde and ketone groups with thiol and amine groups in proteins and DNA^30,31^. The aldehyde and ketone groups are also known to react with oxygen, producing reactive oxygen species^32,33^. While sugars can exist in both cyclic (furanose and pyranose) and non-cyclic forms, only the non-cyclic forms exhibit biotoxicity. Since C5-6 sugars predominantly exist in cyclic forms, with the proportion of non-cyclic forms being less than 0.1% in solution^34^, their biotoxicity are negligible. On the other hand, tetroses do not form cyclic structures except for C4a, and even C4a exhibits a significant proportion of its non-cyclic form in solution (>10%)^35^. The difficulty of tetroses in forming cyclic structures would contribute to their relatively high biotoxicity.

Only a few reports on the inhibitory effects of tetroses on microorganisms were published in the 1960s and 1970s. Those studies reported that the growth of *Vibrio cholerae*^36^ and *Lactobacillus* spp.^37^ is completely inhibited by approximately 3 mM (0.36 g/L) and 20-60 mM (2.4-7.2 g/L) of D-ery, respectively. There have been no reports on the inhibition of bacterial growth by C4k. The present study is the first to systematically investigate the inhibitory effects of tetroses on phylogenetically diverse bacteria. The finding that the inhibitory effect of L-eru, which does not form a ring structure, is greater than that of D-ery aligns with the above-mentioned inhibitory mechanism.

The tetrose-utilizing bacteria isolated in this study had a higher resistance to D-ery and L-eru than the model bacterial strains (Figure 2I-L). One possible factor contributing to the resistance could be the reduction of intracellular concentrations of D-ery and L-eru due to their catabolic capabilities. However, this does not explain the high L-eru resistance observed in strains Ery-7 and Ery-28b (Figure 2I and J), which lack the capacity to catabolize L-eru (Figure 2E and F). Some microorganisms may be able to assimilate tetroses as the carbon source, but not the energy source, as inferred from the observation that low concentrations of tetroses promoted the growth of *C. glutamicum* (Figure 1D). It is conceivable that such anabolic metabolisms may contribute to the tolerance, although further validation is needed. The isolates obtained in this study are valuable for exploring the molecular mechanisms of microbial tetrose resistance, which will serve as fundamental information for biomanufacturing processes utilizing the abiotically synthesized sugars.

We successfully isolated phylogenetically diverse bacteria capable of growing on tetroses (Table 1 and Figure 2A-H). There are only a few reports in the 1950s on bacteria growing on tetroses; *Alcaligenes faecalis* and *Aerobacter aerogenes* were reported to grow on D-ery and L-eru^38,39^. The primary reason why microbial utilization of tetroses has not been extensively studied is likely due to their rarity in nature, resulting in a lack of motivation to investigate it. This study is the first to discover tetrose-utilizing bacteria from natural environments, driven by an unprecedented motivation to improve the efficiency of biomanufacturing based on the abiotically synthesized sugars. Among the isolates, strain Eru-33a was able to grow even in the media supplemented with 4 g/L of D-ery or L-eru, concentrations that completely inhibit the growth of other bacteria. Understanding the molecular mechanisms of tetrose metabolisms in these isolates is expected to contribute to the effective use of the abiotically synthesized sugars for biomanufacturing.

The uptake and catabolic pathways of tetroses in microorganisms remain to be elucidated. On the other hand, since C4 sugar alcohols, in particular erythritol, are used worldwide as sweeteners^40^, not only their biosynthetic pathways^41^ but also their catabolic pathways have been investigated. Two distinct catabolic pathways for C4 sugar alcohols have been identified in different bacterial phylogenetic groups. The α-proteobacteria strains, including *Brucella* spp. and *Ochrobactrum* spp., produce D-erythrose-4-phosphate (D-ery-4P) from erythritol through a five-step reaction, with D/L-erythrulose-1-phosphate and -4-phosphate (D/L-eru-1P and -4P) as intermediates^42-44^ (Figure S3A). On the other hand, *Mycolicibacterium smegmatis*, belonging to Actinomycetota, produces D-ery-4P from erythritol and D/L-threitol through three- or four-step reactions with D/L-eru and D/L-eru-4P as intermediates^45^ (Figure S3B-D). In both catabolic pathways for C4 sugar alcohols, D-ery-4P serves as an intermediate product of the pentose phosphate pathway, subsequently entering other primary metabolic pathways (e.g., glycolysis). We speculate that the isolates obtained in this study possess phosphotransferase systems (PTSs) that simultaneously perform uptake and phosphorylation, and/or sets of transporters and kinases specific to each tetrose. The isolates could catabolize tetroses by using those uptake systems and finally producing the hub metabolite, D-ery-4P. We are currently conducting genomic and transcriptomic analyses of the isolates to identify unknown uptake apparatus for tetroses, in addition to elucidating their catabolic pathways.

The isolates obtained in this study were able to grow in the media supplemented with 5% of the abiotically synthesized sugars, which significantly inhibit growth of typical bacteria. Additionally, these isolates were capable of consuming tetroses present in the abiotically synthesized sugars (Figure 3A-D). In particular, strain Eru-33a was able to consume not only C4a and C4k but also C1-3 compounds present in the abiotically synthesized sugars (Figure 3D and S2D). Although the concentrations of C1-3 compounds are lower than those of tetroses, it may be necessary to use microorganisms capable of metabolizing C1-3 compounds to increase the utilization efficiency of the abiotically synthesized sugars. In particular, strain Eru-33a, with its superior C1-4 consumption activity and resistance to tetroses, is expected to be an excellent model organism and a valuable source of genes and enzymes for the study of effective utilization of the abiotically synthesized sugars.

## CONCLUSION

In this study, we successfully isolated bacteria capable of growing by catabolizing tetroses, a microbial metabolism rarely reported to date. The tetrose-utilizing isolates exhibited higher resistance to tetroses compared to model bacterial strains. Notably, strain Eru-33a was able to grow in high concentrations of the abiotically synthesized sugars, which typically inhibit bacterial growth, and could consume the tetroses contained therein. Elucidating the molecular mechanisms of tetrose catabolism and tolerance of these isolates will provide valuable insights into the development of biomanufacturing processes using the abiotically synthesized sugars as feedstocks.

## ASSOCIATED CONTENT

### Data Availability Statement

The original contributions presented in the study are included in the article/Supporting Information. The nucleotide sequence data obtained from the isolates in the present study have been submitted to the DNA Data Bank of Japan (DDBJ) under accession numbers LC847118 to LC847125.

### Supporting Information

The Supporting Information is available free of charge at XXX. Supplementary Figures S1-S3 (PDF)

## Author Contributions

Conceptualization, S.N. and S.K.; data curation, S.N. and S.K.; formal analysis, K.K., and S.K.; funding acquisition, S.N. and S.K.; investigation, K.K., K.I., H.T., and H.N.; methodology, H.T. and S.K.; resources, K.K., H.T., and H.N.; visualization, S.K..; writing–original draft preparation, S.K.; and writing–review and editing, K.K, I.K., H.T., H.N., S.N., and S.K.

## Notes

The authors declare no competing financial interest.

## ACKNOWLEDGMENT

This study funded by JST-Mirai Program Grant No. JPMJMI22E5, Japan. The authors would like to express appreciation to Ms. Ai Miura, Ms. Mika Yamamoto, Ms. Keiko Fujikawa and Ms. Sharbanee Mitra for technical assistance.

## ABBREVIATIONS

ery: erythrose
eru: erythrulose
glc: glucose
fru: fructose
C1: one carbon aldose (formaldehyde)
C2: two-carbon aldose (glycolaldehyde)
C3a: three-carbon aldose (glyceraldehyde)
C3k: three-carbon-ketose (dihydroxyacetone)
C4a: four-carbon aldoses
C4k: four-carbon ketoses

