## Supplemental Figures S1-S3 for "Isolation of bacteria that catabolize abiotically synthesized tetroses via the formose reaction"

[1] Bioproduction Research Institute, National Institute of Advanced Industrial Science and Technology, 2-17-2-1, Tsukisamu-Higashi, Toyohira, Sapporo, 062-8517 Japan; [2] Research Center for Solar Energy Chemistry, Graduate School of Engineering Science, Osaka University, 1-3 Machikaneyama, Toyonaka, Osaka, 560-8531 Japan; [3] Presidential Endowed Chair for Platinum Society, The University of Tokyo, 7-3-1 Hongo, Bunkyo-Ku, Tokyo, 113-8656 Japan; [4] Innovative Catalysis Science Division, Institute for Open and Transdisciplinary Research Initiatives (ICS-OTRI), Osaka University, Suita, Osaka, 565-0871 Japan; [5] Division of Applied Bioscience, Graduate School of Agriculture, Hokkaido University, Kita-9 Nishi-9, Kita-ku, Sapporo, 060-8589 Japan.

***Corresponding author:**

Bioproduction Research Institute, National Institute of Advanced Industrial Science and Technology, 2-17-2-1, Tsukisamu-Higashi, Toyohira, Sapporo, 062-8517 Japan

Document prepared:

Supplementary materials: Figures S1-S3

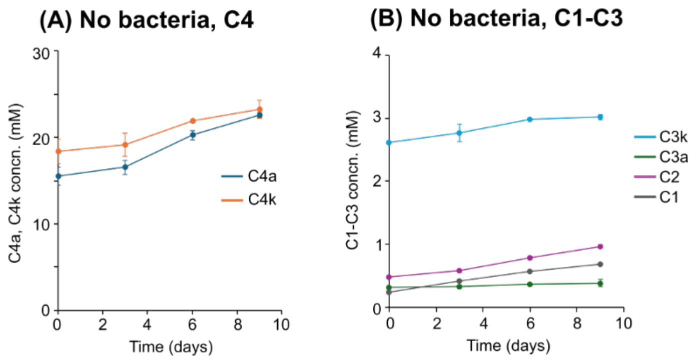


**Figure S1.** Concentrations of C4 sugars (A) and C1-C3 compounds (B) under the uninoculated control. The minimal medium supplemented with 5% of the abiotically synthesized sugars were incubated at 30°C with agitation (180 rpm). Data are presented as the means of three independent cultures. Error bars represent standard deviations.

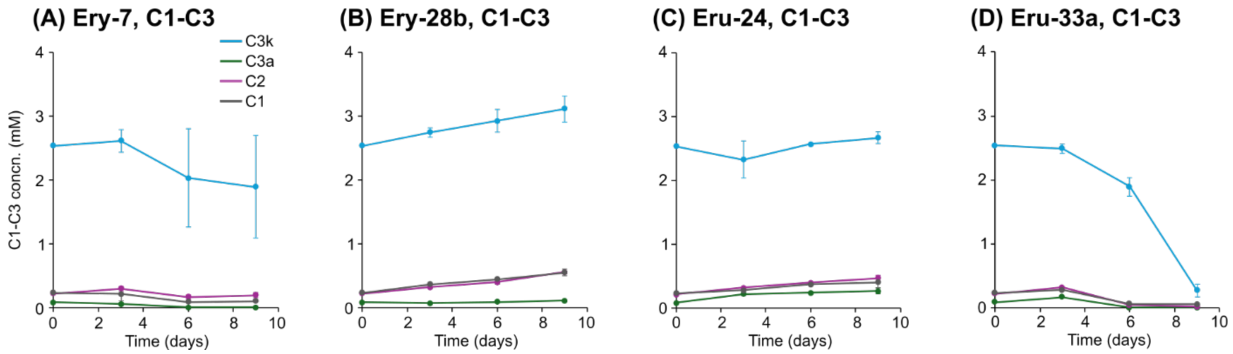


**Figure S2.** Consumption of C1-3 compounds in the abiotically synthesized sugars by the tetrose-utilizing isolates. Each isolate was cultured in the minimal medium supplemented with 5% of the abiotically synthesized sugars and the changes in the concentrations of each sugar species were determined. Data are presented as the means of three independent cultures. Error bars represent standard deviations.

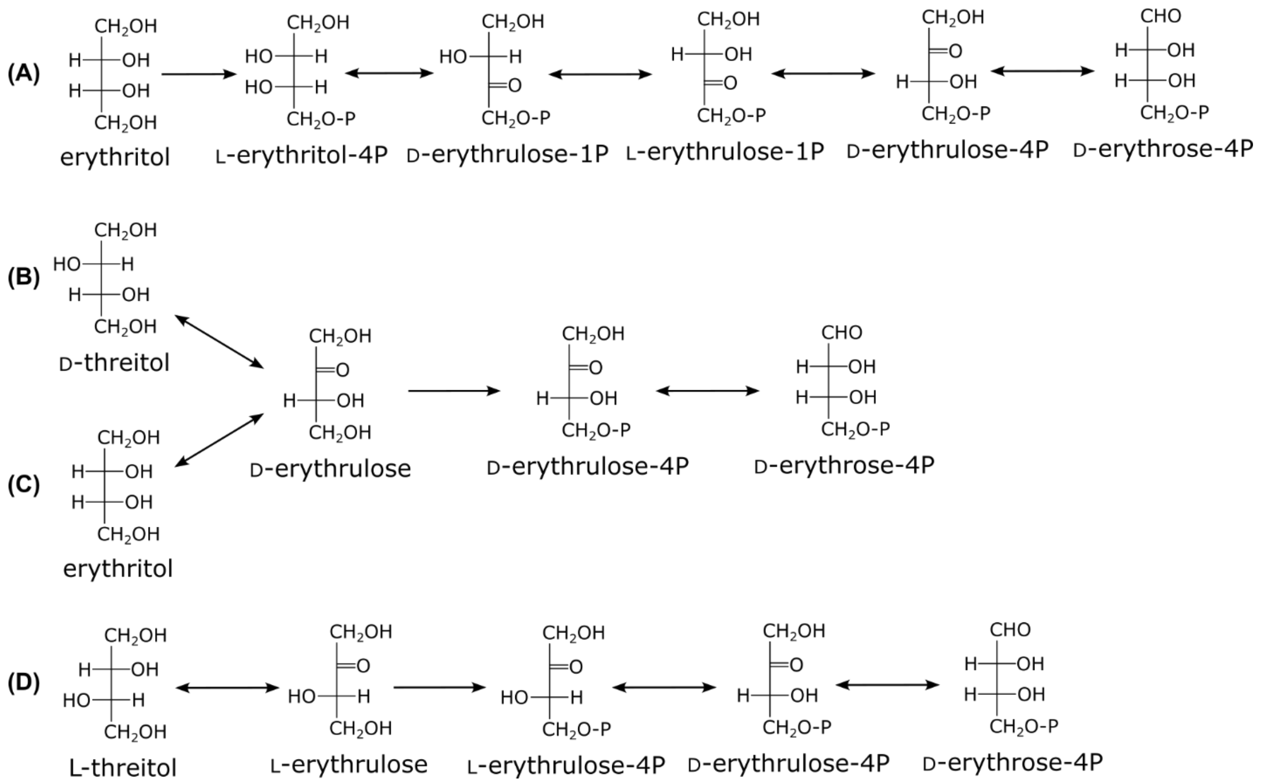


**Figure S3.** Catabolic pathways for C4 sugar alcohols identified in different bacterial strains to date. (A) An erythritol catabolic pathway identified in *Brucella abortus*[43]. (B-D) Catabolic pathways for D-threitol, erythritol, and L-threitol, respectively, identified in *Mycolicibacterium smegmatis*[45].
